# Genetic Mapping in Autohexaploid Sweet Potato with Low-coverage NGS-based Genotyping Data

**DOI:** 10.1101/789198

**Authors:** Eiji Yamamoto, Kenta Shirasawa, Takumi Kimura, Yuki Monden, Masaru Tanaka, Sachiko Isobe

## Abstract

Next-generation sequencing (NGS)-based genotyping methods can generate numerous genetic markers in a single experiment and have contributed to plant genetic mapping. However, the benefits of NGS-based methods are limited in autopolyploids as their genetic segregation mode is complex. Moreover, autopolyploids have large genomes and require abundant sequencing data to obtain sufficient genetic markers. There are several methods for genetic mapping in autopolyploids. These approaches may be impractical for plant genetic studies as they require large amounts of data and are not cost-effective. In the present study, we propose a simple strategy for genetic mapping of polyploids in a cost-effective manner. The allele dosage probabilities calculated from NGS read counts were used in association analyses to detect loci associated with specific phenotypes. This approach is superior to conventional methods of determining allele dosage, which usually result in the filtering of many genetic markers with low read depth. The validity of the strategy was demonstrated using real phenotype data from autohexaploid sweet potato populations to detect genetic loci for both qualitative and quantitative traits, the latter of which required the use of allele dosage probabilities for the detection of loci. We demonstrate that this proposed method is useful with reasonable NGS read counts.

## Introduction

Recent advances in next-generation sequencing (NGS) technology have revolutionized genomics-assisted breeding. NGS-based genotyping by sequencing (GBS; Elshire *et al*. 2011) and restriction site-associated DNA sequencing (RAD-seq; Baird *et al*. 2008) have enabled the development of numerous genetic markers in a single experiment (Kumar *et al*. 2012). They have been used to construct high-density genetic linkage maps (Poland and Rife 2012) and genetic maps of agronomically important traits. These technologies are highly effective with diploid species but present numerous application challenges with autopolyploid species (Bourke *et al*. 2018).

Polyploidy is the presence of multiple sets of chromosomes in a single plant and is a common occurrence in the plant kingdom. Polyploid plant species are often valuable as crops, as their genome multiplication results in comparatively larger yields. In addition, polyploidy often leads to heterosis, gene redundancy, loss of self-incompatibility, and gains in asexual reproduction (Comai 2005). In allopolyploid species, such as cotton and wheat, preferential paring dictates meiotic chromosome behavior similar to diploids. As this mechanism resembles that seen in diploids, currently available genetic approaches can be readily applied to allopolyploids. By contrast, autopolyploids have several different genotypes per locus. Consequently, the existing approaches designed for diploids are not applicable to autopolyploids (Bourke *et al*. 2018). A possible solution for this problem is the use of Mendelian markers such as Simplex × Nullplex (SN) and Simplex × Simplex (SS). The mode of inheritance of Mendelian markers resembles that for the genetic markers in diploid species. Thus, they apply to the theories and/or tools developed for diploids (Tennessen *et al*. 2014; Vukosavljev *et al*. 2016; Shirasawa *et al*. 2017). To detect genetic loci with simple inheritance and/or a high proportion of variance explained, the use of Mendelian markers alone may suffice. However, genetic mapping based on allele dosage information may be required for more complex phenotypes (Rosyara *et al*. 2016).

To use multiple-dose markers, the allele dosage must be determined. Several techniques can be used to estimate allele dosage in polyploids (Serang *et al*. 2012; Gerard *et al*. 2018; Wadl *et al*. 2018). These techniques have enabled the development of genetic mapping methods for polyploids (Rosyara *et al*. 2016; da Silva Pereira *et al*. 2019). Even with the available tools, accurate allele dosage estimation demands an adequate amount of high-quality data. To meet this requirement, the first allele dosage estimation method was developed for SNP-chip data (Serang *et al*. 2012). For NGS-based genotyping, abundant sequence data are needed for species at higher ploidy levels and with larger genome sizes. Gerard *et al*. (2018) recommended read depths > 25 and >90 to obtain accurate allele dosages for autotetraploids and autohexaploids, respectively. Wadl *et al*. (2018) developed a GBS pipeline for polyploid study. They reported that >100 reads were necessary to achieve 95% accuracy for allele dosage estimation in autohexaploid species. However, this approach may be impractical for plant genetic studies as it is not cost-effective.

In the present study, we propose a genetic mapping strategy for autopolyploids. To validate the proposed strategy, we used two genetic mapping populations in sweet potato (*Ipomoea batatas* (L.) Lam. Sweet potato is a hexaploid species with 90 chromosomes (2n = 6x = 90), and the most possible genome structure is considered B1B1B2B2B2B2 (Shiotani and Kawase 1989). The main objective of this study was to perform genetic mapping in polyploids in a cost-effective manner (i.e., with low-coverage NGS-based genotyping data). In our proposed method, the allele dosage probability for each single-nucleotide polymorphism (SNP) marker site is calculated on the basis of read count information from low-coverage double digest (dd) RAD-seq genotyping data. We do not attempt to determine allele dosage where the counts are too small. Alternatively, allele dosage probabilities can be used in subsequent genetic mapping. This idea is similar to a previous study that used continuous genotype values from signal intensity of SNP-chips for genetic mapping (Grandke *et al*. 2016). In the present study, we applied this idea to low-coverage NGS-based genotyping data. In this manner, the maximum use of the available genetic marker information can be made. Our strategy successfully detected genetic loci for sweet potato agronomic traits.

## MATERIALS AND METHODS

### Plant materials and phenotypes

Two populations of autohexaploid sweet potato (2n = 6x = 90) were used. One was the F1 derived from reciprocal crosses between the major Chinese variety Xushu 18 and the wild sweet potato (*Ipomoea trifida*) (K123–11); hereafter, this population is referred to as KX-F1. The other population originated from self-pollinated (S1) Xushu 18 (n = 248) used in a previous study (Shirasawa *et al*. 2017); hereafter, this population is called X18-S1. These materials were developed by the Kyushu Okinawa Agricultural Research Center of the National Agricultural and Food Research Organization (KARC/NARO). KX-F1 was phenotyped for color and internode length and X18-S1 was phenotyped for color (Table 1). X18-S1 was planted in a field at Okayama University from June through November 2016. KX-F1 was planted in a field at the Miyakonojo Research Station of KARC/NARO from June through November 2016.

**Table 1.**
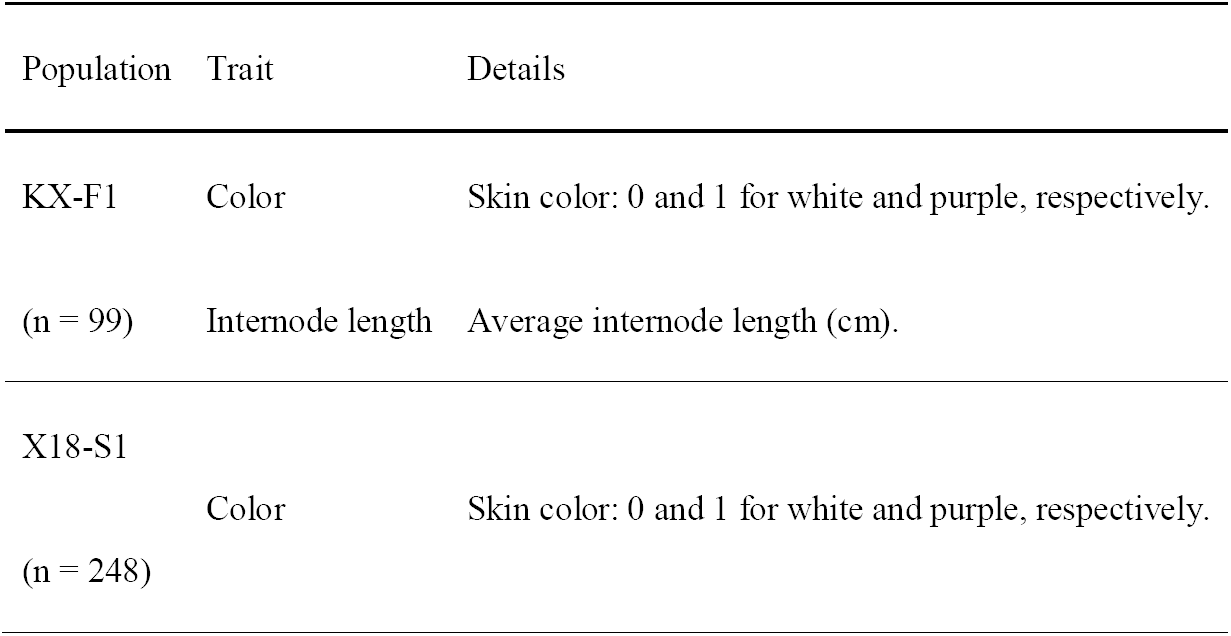
Traits analyzed.

### ddRAD-seq and variant calling

The genomic DNA of KX-F1 was extracted from the leaves with a DNeasy Plant Mini Kit (Qiagen, Hilden, Germany). The ddRAD-seq analyses were performed as described in Shirasawa *et al*. (2016), and ddRAD-Seq libraries were constructed using the restriction enzymes *Pst*I and *Msp*I. DNA fragments of 300–900 bp in length were fractionated using BluePippin (Sage Science, Beverly, MA, USA). The nucleotide sequences of the libraries were determined on the HiSeq 2000 and HiSeq 4000 platforms (Illumina, San Diego, CA, USA) in paired-end mode (93 base or 101 base). The ddRAD-seq reads for the X18-S1 populations were obtained from the DNA Data Bank of Japan (DDBJ) sequence archive under accession numbers DRA004836, DRA004837, and DRA004838.

Data were processed as described in Shirasawa *et al*. (2017). Low-quality sequences were removed and adapters were trimmed with PRINSEQ v. 0.20.4 (Schmieder and Edwards 2011) and fastx_clipper in the FASTX-Toolkit v. 0.0.13 (http://hannonlab.cshl.edu/fastx_toolkit). The filtered reads were mapped onto the *I*. *trifida* “Mx23Hm” (ITR_r1.0) genome sequence (Hirakawa *et al*. 2015) using Bowtie 2 v. 2.2.3 (Langmead and Salzberg 2012). The parameters were set as maximum fragment size length (I) = 1000 and the ‘–sensitive’ preset of the ‘–end-to-end’ mode. The sequence alignment/map (SAM) files were converted into binary sequence alignment/map (BAM) files and subjected to SNP calling using the mpileup option in SAMtools v. 0.1.19 (Li *et al*. 2009) and the mpileup2snp option of VarScan 2 v. 2.3 (Koboldt *et al*. 2012). The output was a VCF file with SNP data. Information in the VCF files was loaded into the R platform via “read.vcfR” in *vcfR* (Knaus and Grunwald 2017). Markers with missing values > 0.5 were filtered out using an in-house R script.

### 𝒳^2^ test to search for Mendelian markers

For autopolyploid species, SN and SS were selected as they reflect a simple Mendelian inheritance mode. To search for potential Mendelian markers, the **𝒳**^2^ test was applied to identify deviation from the expected ratios. Information in the genotype field (GT) of the VCF files was used as the input genotype data. The **𝒳**^2^ test was performed using ‘chisq.test’ in R (R Core Team 2018).

### Allele dosage estimation

Here, the allele dosage was the dosage of the reference genome type allele for each SNP locus. For allele dosage estimation, information on the depths of the total reads (DP) and the reference type reads (RD) were extracted from the VCF files using ‘extract.gt’ in *vcfR* of R (Knaus and Grunwald 2017). DP < 10 and DP > 1000 were filtered out. For N-ploid species, the possible allele dosage states are *d* □ {0/N, 1/N, …, N/N}. For real data, errors in the experimental procedure introduce bias relative to the theoretical probabilities. Therefore, the expected value of the allele dosage *d* in the real data was determined as *r* □ {0/N+e, 1/N, …, N/N-e} where e is the unknown error probability. For the error probability, 0.001 was used. For a given DP and RD, the probability (Pr) of dosage *d*_*i*_ was calculated using the binomial distribution function:

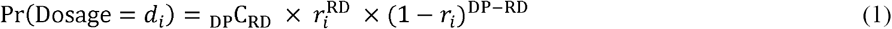

Thus, N + 1 probability values were calculated for each individual at each SNP site. The relative probability (RPr) for the allele dosage *d*_*i*_ was calculated as follows:

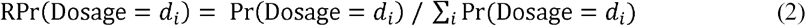

When a value satisfying RPr (Dosage = *d*_*i*=*x*_) was >0.95, RPr (Dosage = *d*_*i*=*x*_) was set to 1 and RPr (Dosage = *d*_*i*_≠*x*) was set to 0. In this way, a matrix *M* was obtained for each SNP marker with individuals as row elements. The column elements were the relative probabilities of the reference type allele dosages calculated by equation (2). Calculation with the binomial distribution function was performed in ‘dbinom’ in R (R Core Team 2018). The SNP markers included potential monomorphic markers. These were identified using a major genotype frequency, namely, the ratio of individuals with a specific allele dosage at a given SNP marker. The major genotype frequency was estimated by aggregating the column elements of the matrix *M* developed from equations (1) and (2).

### Association analyses

The association between marker genotype and phenotype was tested with a generalized linear model (GLM) using

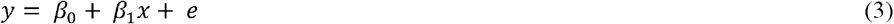

for continuous traits, and

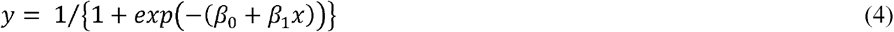

for binary traits.

The term *y* is a vector of phenotypic values, *β*_0_ is the intercept, *β*_1_ is a vector for the effects of each allele dosage state, *e* is the error, and *x* is the genotype information on each SNP marker.

The statistical significance of each SNP marker was assessed using the F-test:

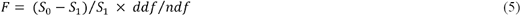

where *S*_0_ and *S*_1_ are residual sums of squares for phenotypic values and the error term in the fitted GLM, respectively. The terms *ndf* and *ddf* are the degrees of freedom of the phenotypic values and rank of *x* in equation (3) or (4).

GLM fitting was performed using ‘glm’ in R (R Core Team 2018). The augment family functions “binomial” and “gaussian” were used for binary and continuous traits, respectively. The genome-wide significant threshold was determined based on a Bonferroni correction. To build Manhattan plots of the association analyses, the SNP markers were allocated to 15 homologous linkage groups identified in a previous study (Shirasawa *et al*. 2017). Note that the order of the SNP markers in a homologous linkage group did not correspond to a genetic or physical map position because the genetic map of Shirasawa *et al*. (2017) did not include SNP marker information except for the SS markers and the physical map of the reference genome (Hirakawa *et al*. 2015) used in the present study was fragmented and did not represent all of the chromosome sequences. The Manhattan and quantile–quantile (QQ) plots were created with ‘manhattan’ and ‘qq,’ respectively, in *qqman* of R (Turner 2014).

### Data availability

The phenotype and genotype data, the R scripts, and the R library developed for the present study are available at https://github.com/RMcj625fE4/polyploidNgsMapping. The sequence data from the ddRAD-seq libraries are available in the DDBJ sequence read archive under accession numbers DRA008654–DRA008655 and DRA004836–DRA004838 for KX-F1 and X18-S1, respectively.

## RESULTS

### Importance of allele dosage estimation

Mendelian markers such as SN and SS are useful for genetic analyses in autopolyploid species. We estimated the number of possible SN or SS markers from the genotype field information in VCF files. Hereafter, we refer to this as a “diploidized genotype” (Rosyara *et al*. 2016). Table 2 shows the potential SN and SS markers. The indicated values are markers that were not rejected by the *χ*^2^ test within the *p*-values. Therefore, more severe conditions were used to select possible SN and/or SS markers in practical genetic studies (Shirasawa *et al*. 2017). At least 43.5% and 44.2% of the SNP markers were discarded as multiplex markers from KX-F1 and X18-S1, respectively. Thus, Mendelian marker (SN and SS) selection ignores substantial available multiplex marker information.

**Table 2.**
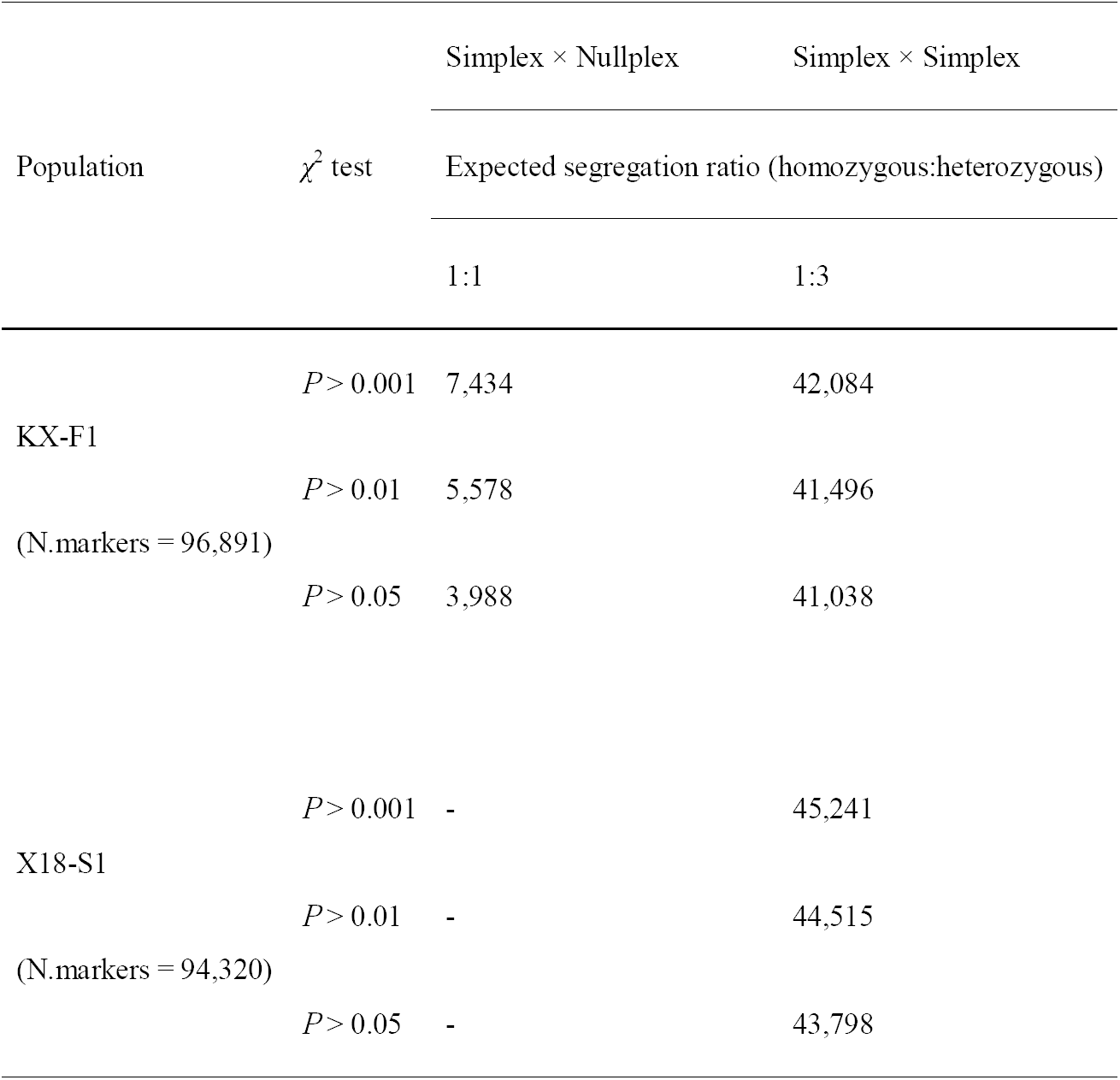
Markers expected to follow Mendelian segregation.

### Allele dosage estimation

For NGS genotyping of autopolyploid species, the read depth at the SNP marker site is critical for accurate allele dosage estimation (Gerard *et al*. 2018; Wadl *et al*. 2018). Read depth and allele dosage estimation accuracy for all SNP marker sites of KX-F1 and X18-S1 are summarized in Figure 1A and B. Here, we defined allele dosage accuracy as the maximum value in an allele dosage probability array at an individual SNP. For SNP marker Itr_sc000001.1_24872 of the individual KX001, the allele dosage accuracy was 0.74 (Fig. 1C). There were wide differences in read depth among individuals even within a single SNP marker. These discrepancies cause large variation in accuracy (Fig. 1A and B).

**Fig. 1.**
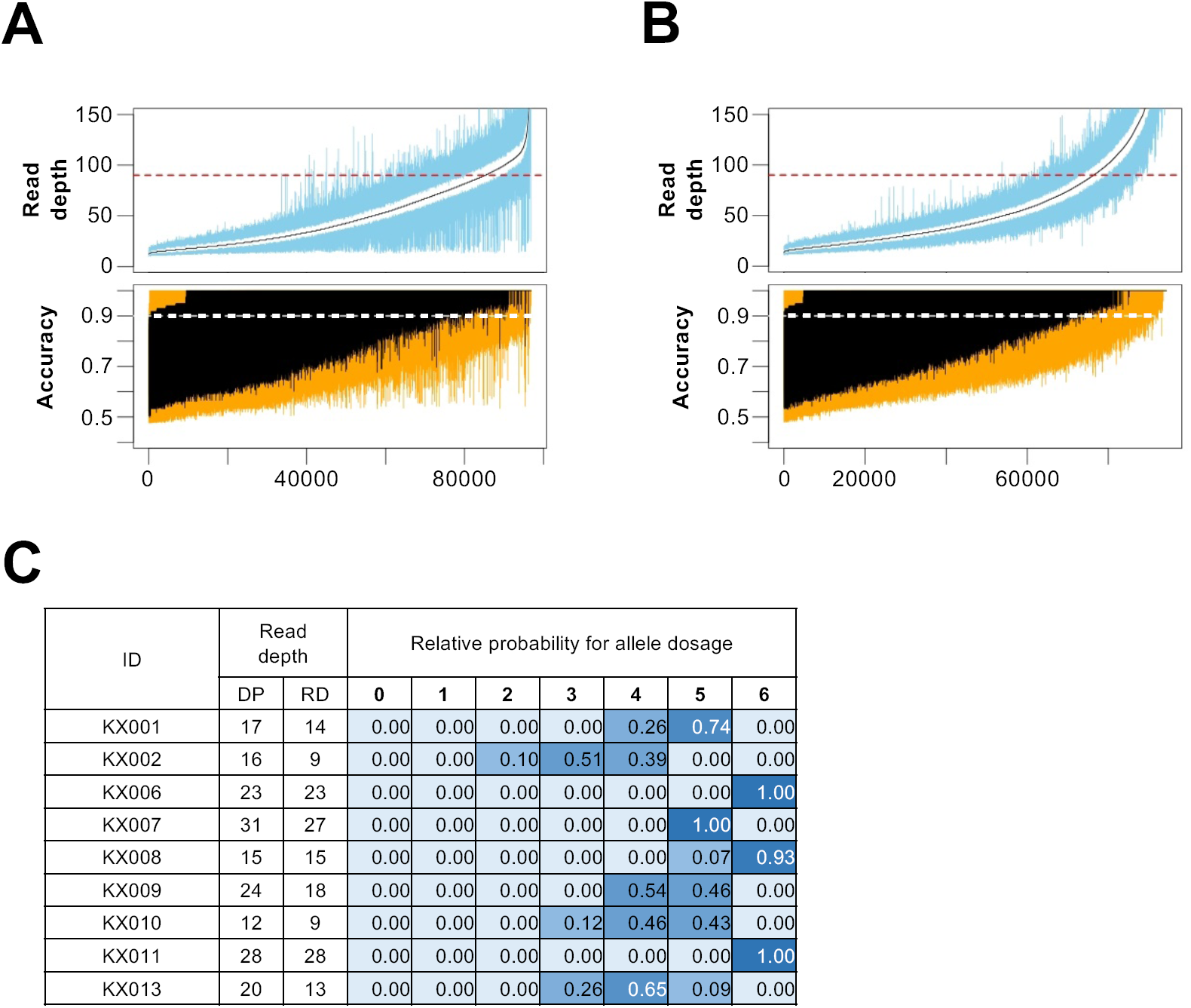
Allele dosage estimation. (A and B) Relationship between read depth (upper panels) and accuracy of estimated allele dosage (lower panels) for KX-F1 (A) and X18-S1 (B). Accuracy of the estimated allele dosage was the maximum value in a relative probability array for allele dosage at a given SNP marker. The *x*-axes indicate SNP markers ordered according to the median of the total read depth. Black lines indicate the median. The upper and lower blue lines in the upper panels indicate the 25th and 75th percentile read depths, respectively. The red dashed lines indicate read depth = 90. The upper and lower orange lines in the lower panels indicate the 25th and 75th percentiles, respectively. The white dashed lines indicate 0.9 accuracy. (C) Example of allele dosage estimation. Part of the result at SNP Itr_sc000001.1_24872 in KX-F1. ID = name of individuals. DP and RD in the read depth column mean the total and reference genome type read counts, respectively. Values in the colored cells indicate the relative allele dosage probabilities.

SNPs with accuracy > 0.9 (white lines in the lower panels of Fig. 1A and B) correspond to SNPs with read depths > 90 (red lines in the upper panels of Fig. 1A and B). This finding was consistent with the conclusions of previous studies (Gerard *et al*. 2018; Wadl *et al*. 2018). When we filter SNP markers on the condition that the median read depth for each individual is >90 (Gerard *et al*. 2018), >87% and >80% of the SNP markers are removed from KX-F1 and X18-S1, respectively (Fig. 1A and B). In the present study, we focused on allele dosage probability rather than allele dosage determination (Fig. 1C). We used a relative probability array for each allele dosage state as the input for subsequent genetic mapping analyses. Seven values (probabilities for allele dosages 0/6, 1/6,…, 6/6) were calculated as genotype information at the individual SNP marker (Fig. 1C). SNP markers with estimated major genotype frequencies > 0.95 were filtered out as they furnished too little information for genetic mapping. We obtained 89,407 (92.3%) and 82,007 (86.9%) SNP markers for KX-F1 and X18-S1, respectively.

### Genetic mapping of the agronomic traits

We performed association analyses for the real phenotypes in KX-F1 and X18-S1 (Table 1 and Fig. 2). Color is a qualitative trait with binary phenotypes. Internode length is quantitative and continuous. Association analyses were performed with diploidized genotypes and allele dosage probabilities (Fig. 3 and Supplemental Fig. S1).

**Fig. 2.**
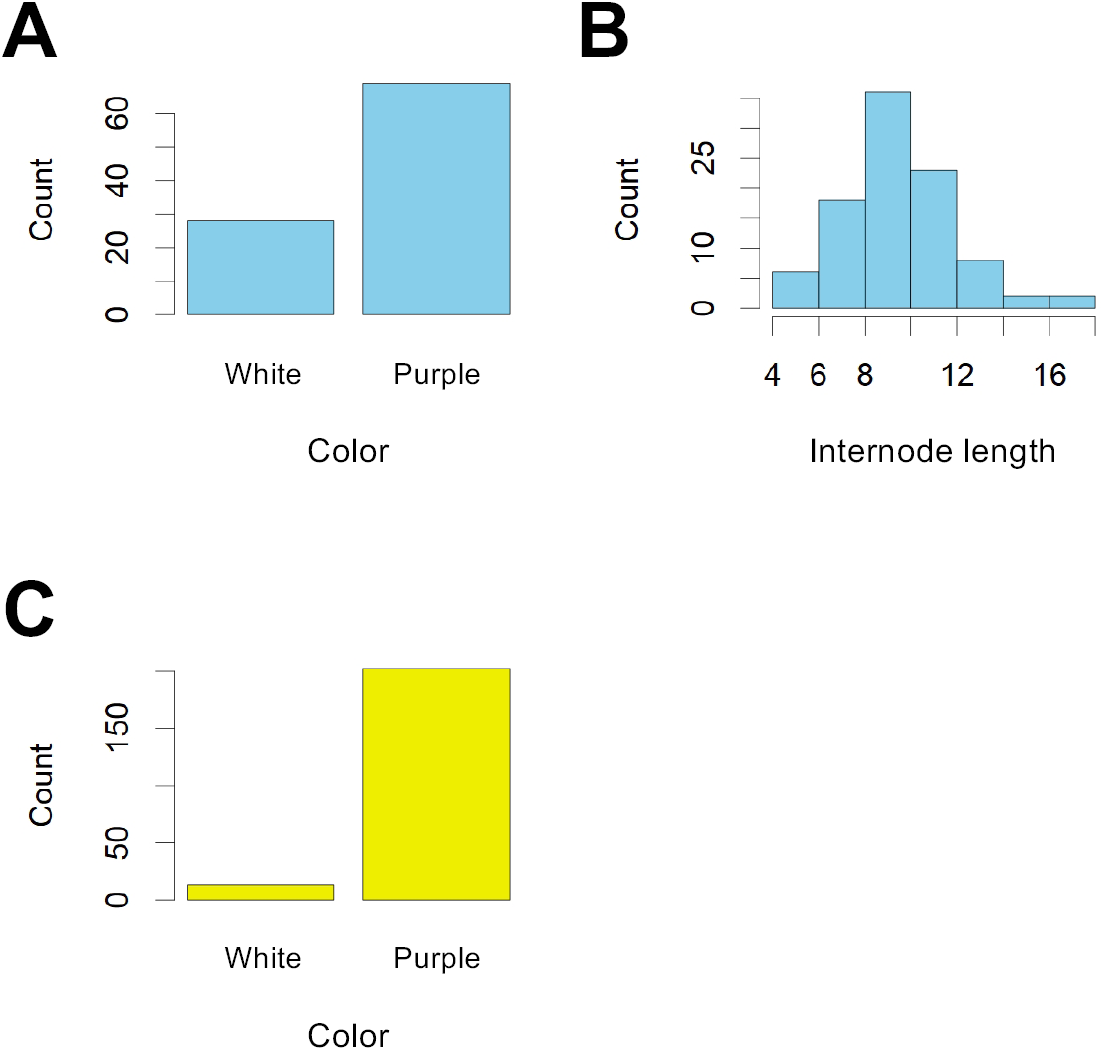
Distribution of phenotypic values. (A) Color of KX-F1. (B) Internode length of KX-F1. (C) Color of X18-S1.

**Fig. 3.**
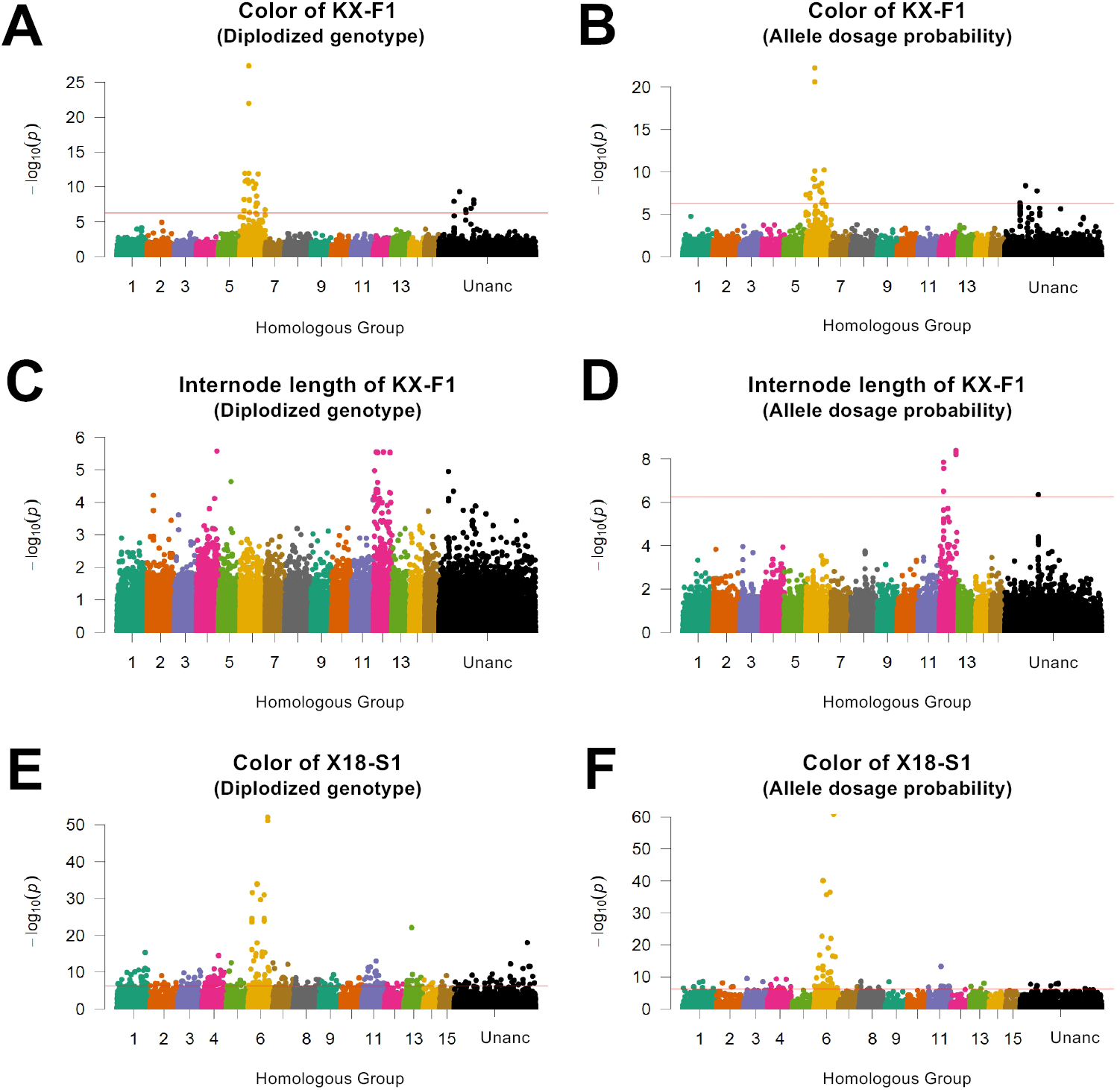
Genetic mapping results. Manhattan plots of the association analyses for the agronomic traits. The red lines in the Manhattan plots indicate a 5% genome-wide significance threshold based on a Bonferroni correction.

For color, the association analyses with diploidized genotypes and allele dosage probabilities were similar for both KX-F1 and X18-S1 (Fig. 3A, B, E, and F). Strong significant peaks (Itr_sc000236.1_59664 and Itr_sc001350.1_30359 for KX-F1 and X18-S1, respectively) were detected on homologous linkage group 6. Comparisons of the phenotypic values and the estimated allele dosages at the significant SNPs indicated that the phenotype inheritance mode was simplex dominant. The homozygote of an allele had no effect, whereas the heterozygote or homozygote of another allele affected the phenotype (Fig. 4A and B). For color, both the diploidized genotype and the allele dosage were applicable for locus detection.

**Fig. 4.**
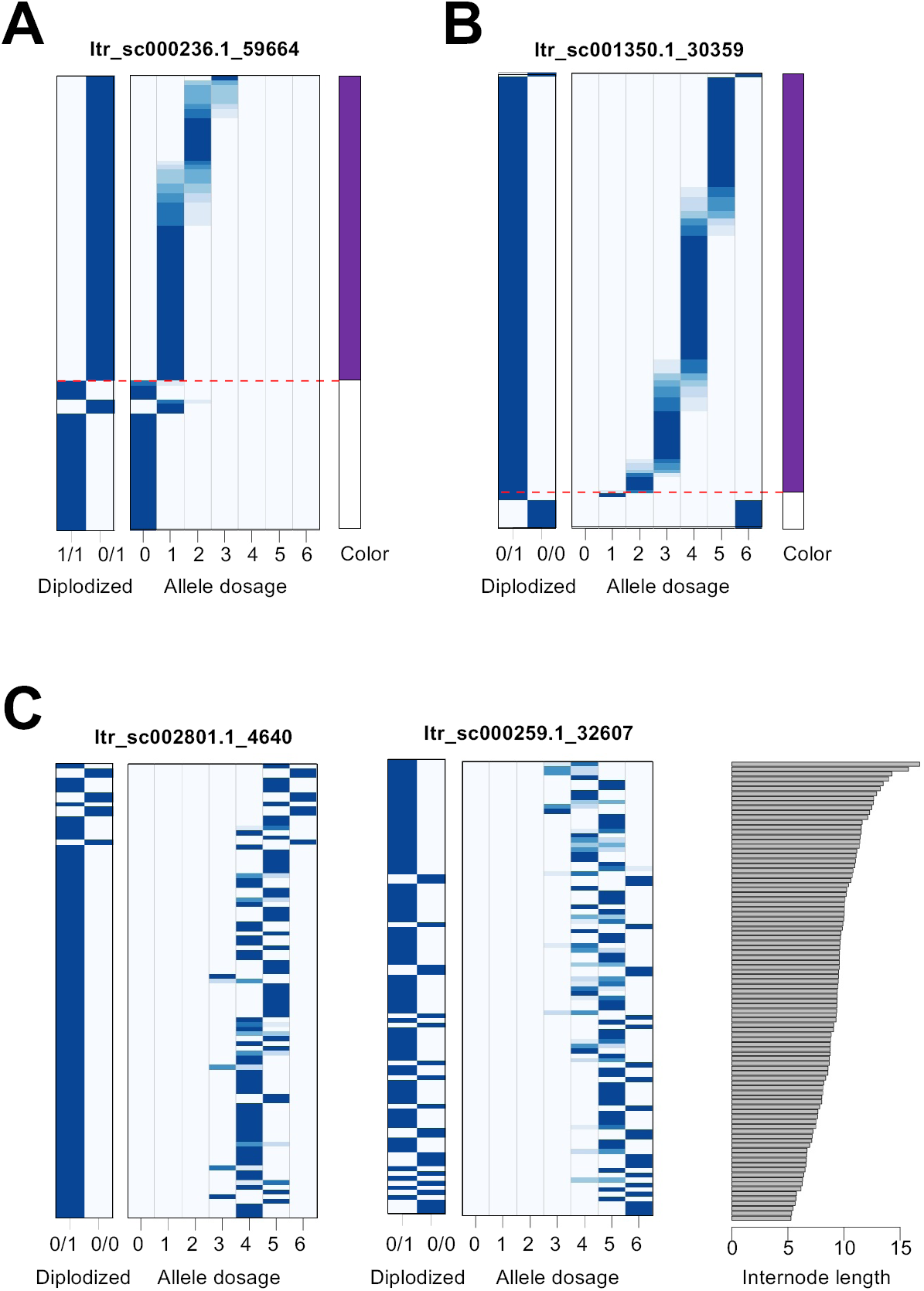
Comparisons between phenotypic values and allele dosage probabilities for the SNP markers. Boxes in each panel indicate allele dosage probabilities. Rows indicate individuals and blue cells in the columns indicate individual allele dosage probabilities. The gradient from white to blue indicates values ranging from 0–1. Plots on the right side of each panel indicate the phenotype. (A) Color of the KX-F1 population. The red dashed line indicates the boundary between purple and white individuals. (B) Color of the X18-S1 population. The red dashed line indicates the boundary between purple and white individuals. (C) Internode length of KX-F1.

For the internode length of KX-F1, no SNP marker was significant in the association analyses with the diploidized genotype (Fig. 3C). However, allele dosage probability detected a significant peak on homologous group 12 (Fig. 3D). Figure 4C compares internode length and allele dosage at the top and the fourth-highest signals. These SNPs showed different genotype patterns. For the top signal (Itr_sc002801.1_4640), the internode length increased with reference allele dosage (Fig. 4C). By contrast, at the fourth-highest peak (Itr_sc000259.1_32607), the internode length decreased as reference allele dosage increased (Fig. 4C). For the diploidized genotype, the dosage 6 column in Figure 4 is defined as a homozygous reference allele (0/0), whereas the others are heterozygous (0/1). For Itr_sc002801.1_4640, all 0/0 (allele dosage 6) individuals presented with long internodes, but the number of individuals was too small to achieve statistical significance (Fig. 4C). For Itr_sc000259.1_32607, allele dosages 3 and 4 presented with longer internodes (Fig. 4C). However, in the diploidized genotype, allele dosages 2–5 were heterozygous (0/1). Thus, it was impossible to establish any association between phenotype and genotype. Allele dosage estimation was therefore necessary to detect the locus for internode length (Fig. 3D).

## DISCUSSION

We genotyped populations using ddRAD-seq. In general, polyploid species have large genomes and multiple allele dosages. Therefore, numerous markers must be developed to capture genetic variation in the entire genomic region. NGS-based genotyping such as RAD-seq and GBS are powerful because they generate hundreds to thousands of SNP markers per experiment. High read counts per SNP marker are necessary for accurate allele dosage estimation (Gerard *et al*. 2018; Wadl *et al*. 2018). However, this approach is often impractical for plant breeding because of the cost. In the present study, the filtration of SNP markers for accurate allele dosage estimation purges most SNP markers (Fig. 1A and B). Therefore, we focused on allele dosage probability rather than determination. Thus, we used a comparatively larger number of SNP markers as “genotype” data in our association analyses.

Genetic mapping of internode length demonstrated the importance of allele dosage estimation. No significant peaks for this trait were detected without allele dosage estimation (Fig. 3C and D). However, in the genetic mapping of color, allele dosage estimation was not required for the detection of significant peaks, as the phenotype inheritance mode was simplex dominant. In this case, genotype × phenotype relationships are either homozygous or heterozygous (Fig. 4A and B). For X18-S1, however, another important aspect of allele dosage estimation was revealed (Fig. 4B). This relates to the maximum use of the available genetic marker information. Mendelian markers such as SS are often used in genetic studies on polyploid species as they behave in the same way as genetic markers for diploid species (Tennessen *et al*. 2014; Vukosavljev *et al*., 2016; Shirasawa *et al*. 2017). Thus, they are exempt from the problems associated with genetic analyses of autopolyploids (Bourke *et al*. 2018). In the S1 population, the expected SS marker segregation ratio is 1:3 for homozygous (dosage 6) to heterozygous (dosages 1–5) (Table 2). The segregation pattern of Itr_sc001350.1_30359 differs from the expected ratio and is removed when we select Mendelian markers (Fig. 4B). As Itr_sc001350.1_30359 showed the highest –log_10_(*P*) value, it is most closely linked to the gene causing the phenotypic variation (Fig. 3E and F). Thus, Mendelian marker selection discards the most important information.

The strategy used in the present study is simple but was efficacious for real data (Figs. 3 and 4). Unlike other tools, the only prerequisite condition is that the genotype data must be obtained by NGS-based methods. For this reason, the approach used in the present study is widely applicable. Nevertheless, a drawback of this characteristic is that it ignores certain information such as the number of effective alleles in the polyploid genome and the precise chromosomal locations of the genes. Complex methods that specifically determine allele dosage are necessary to precisely map the genetic loci (Bourke *et al*. 2018; da Silva Pereira *et al*. 2019; Rosyara *et al*. 2016). To use them, however, an abundance of high-quality genotype data is necessary. Therefore, we recommend the following genetic mapping strategy for autopolyploid crop species: (1) Perform NGS-based genotyping using a reasonable data volume. (2) If positive results are obtained, increase the volume of sequencing data for genotyping and apply the output towards the complex methods. In this way, the agronomic traits can be genetically mapped in a cost- and labor-effective manner.

The strategies proposed in the present and previous studies assume the application for autopolyploids (Bourke *et al*. 2018). However, there are more complicated situations in the genetics of polyploids. For example, the genome of sugarcane is polyploid and aneuploid, including variable ploidy levels within the species and genetic mapping population (Grivet and Arruda 2002). Genetic analyses that deal with this case have not been established. The development of strategies and methods for such cases are the next challenges for genetics in polyploid species.

## Acknowledgments

The authors thank S. Sasamoto, S. Nakayama, H. Tsuruoka, and C. Minami of the Kazusa DNA Research Institute for their technical assistance. We also thank the Technical Support Center (Operation Unit 3) of KARC/NARO for their assistance in cultivating and phenotyping the sweet potato population. This work was supported by the Kazusa DNA Research Institute Foundation and JSPS KAKENHI (Grant No. 15H04441 to MT and SI).

