## Supplementary figures and images for "Genetic Mapping in Autohexaploid Sweet Potato with Low-coverage NGS-based Genotyping Data"

### Supplemental Fig. S1

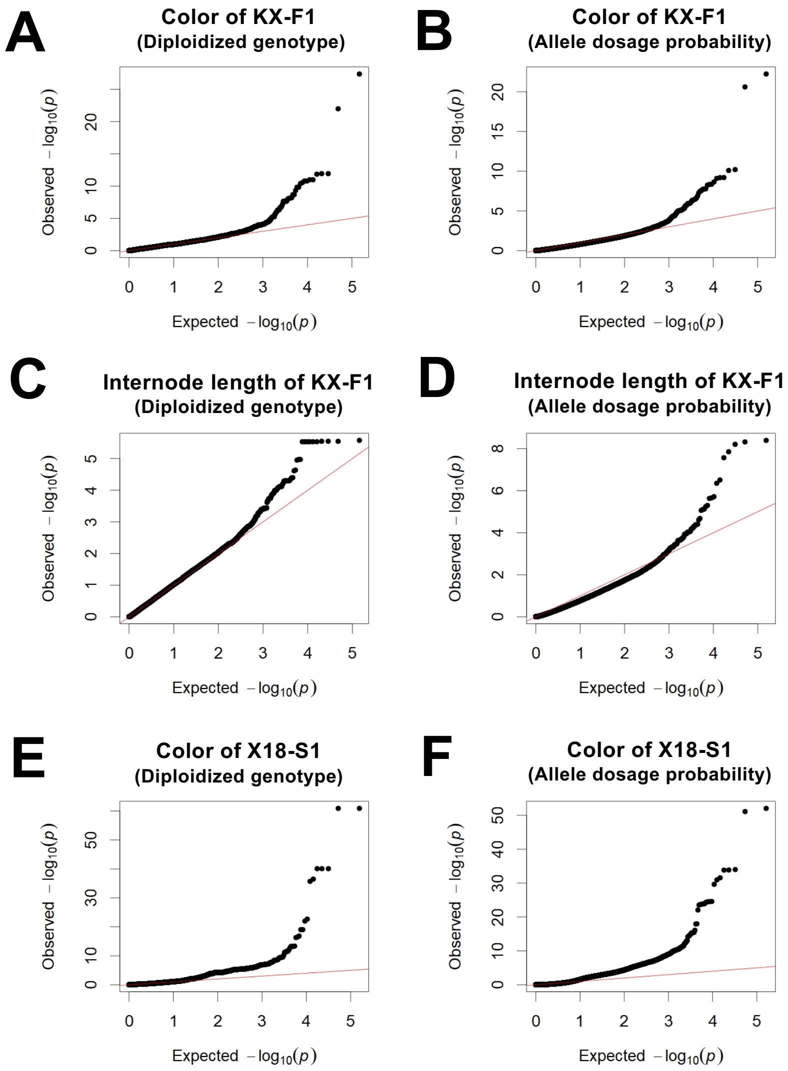


**Figure S1** Quantile–quantile plots for the association analyses.

#### Figure S1
